# Immune Spatial Organization Predicts Metastasis Risk in Aggressive Localized Prostate Cancer

**DOI:** 10.1101/2025.05.15.654324

**Authors:** David D Yang, Aya Abdelnaser, Jeremiah Wala, Alexander J Haas, Alfred A Barney, Eddy Saad, Jett P Crowdis, Cora A Ricker, Jihye Park, Martin T King, Paul L Nguyen, Toni K Choueiri, Alok K Tewari, Keyan Salari, Mary-Ellen Taplin, Chin-Lee Wu, Eliezer M Van Allen

## Abstract

Risk stratification in localized prostate cancer (PCa) remains imprecise. Computational pathology has emerged as an attractive option for improving risk stratification, but current approaches either lack interpretability or focus solely on tumor morphology. Here, we identify, validate, and provide interpretability for an immune microenvironment-derived computational pathology biomarker for high-grade PCa. Using digitized hematoxylin and eosin-stained (H&E) slides from two independent cohorts of patients treated with radical prostatectomy (n=490), we found that spatial clustering of immune clusters, but not immune cell abundance, was independently associated with reduced risk of distant metastasis for high-grade disease but not low-grade. Joint analysis of H&E images and RNA sequencing (n=326) revealed that in high-grade disease, high-cluster samples were enriched in CD8+ T cells, activated memory CD4+ T cells, and Tregs, as well as clonal T cell populations, overall suggestive of an underlying T cell-mediated antitumor response. No such enrichment was found in low-grade tumors. These findings establish immune spatial architecture as a novel, interpretable computational pathology biomarker and provide insight into the immune landscape of high-grade PCa.

## INTRODUCTION

Prostate cancer (PCa) is the second most common cancer in men in the world, with an estimated 1,466,680 new cases in 2022.^1^ While PCa is often indolent and diagnosed at early stages, it is the fifth leading cause of cancer mortality, with an estimated 396,792 deaths in 2022. The clinical management of localized PCa is based on risk stratification, which was first formalized approximately 30 years ago and utilizes clinical and pathologic variables, such as the Gleason score, tumor stage, and prostate-specific antigen (PSA) level at diagnosis, to estimate disease aggressiveness.^2–4^ However, the performance of conventional risk stratification is suboptimal.^5^

In recent years, several novel biomarker approaches to improve risk stratification have emerged. One approach is computational pathology, which utilizes quantitative methods to analyze digitized histopathology slides (typically hematoxylin and eosin [H&E]-stained). With few exceptions,^6^ computational pathology biomarker approaches in PCa fall into two broad categories. In the first group,^7–9^ a whole-slide image is divided into small image tiles, and a deep learning model is trained to learn and extract latent representations (i.e., compressed measurements which capture the most informative features) of each image tile, which are then aggregated into patient-level predictions. This approach has demonstrated strong performance gains compared to conventional risk stratification^10^ but inherently lacks interpretability. The second approach focuses on the tumor region, in essence, refining identification of Gleason patterns and other relevant tumor morphologies, such as cribriform pattern and intraductal carcinoma.^11–13^ While more closely linked to human-interpretable features, this approach ignores non-tumor regions, which contain important prognostic information.^14^ Indeed, emerging evidence suggests that a subpopulation of localized PCa is immune-enriched,^15,16^ particularly high-grade disease^17,18^ and in patients of African ancestry,^19,20^ although these microenvironmental findings have unclear clinical implications in part due to the difficulties in reproducibly measuring these properties from histopathology images.^21,22^ Furthermore, PCa computational pathology biomarkers have thus far largely focused on biochemical recurrence as the clinical endpoint of interest. However, biochemical recurrence is a suboptimal clinical endpoint in localized PCa; it is not a surrogate for overall survival,^23^ as many patients with biochemically recurrent PCa ultimately die of other causes due to the long natural history of the disease and advanced patient age at time of recurrence. Whereas many localized PCa images exist for deriving Gleason scoring models,^13,24–28^ the scarcity of available clinically annotated histopathology data in surgically treated PCa with sufficient duration of clinical follow-up to allow for observation of more relevant endpoints have restricted innovation in understanding additional features that may contribute to distinct disease trajectories.

Here, we aimed to identify a biologically informed computational pathology biomarker in localized PCa that could augment existing tumor-intrinsic Gleason scoring models and improve risk stratification. Specifically, we examined whether reproducible inference of immune features from the prostate tumor microenvironment could be performed using computational pathology strategies. We hypothesized that certain patients with high-grade disease have systemic immune-mediated antitumor responses associated with decreased risk of distant metastasis and that quantifying histologic immune features may allow for the identification of these patients.

## RESULTS

### Study cohorts

We retrospectively identified three cohorts of patients with nonmetastatic PCa who underwent radical prostatectomy (RP) for their disease with digitized H&E slides from the surgical specimen. The Discovery Cohort consisted of 272 patients,^29^ and the independent Validation Cohort consisted of 218 patients. Key baseline characteristics are listed in Table 1. 14% (n=37) of the Discovery Cohort and 17% (n=38) of the Validation Cohort had Gleason 8-10 disease from the prostatectomy specimen. Median follow-up periods after RP were 12.6 and 8.1 years, respectively. These two cohorts allowed for the identification of digital pathology immune features which are associated with the primary endpoint of time to distant metastasis (DM), chosen given the hypothesized role of immune-mediated systemic antitumor activity in select patients. Additionally, we identified 326 patients from The Cancer Genome Atlas (TCGA) who underwent RP for PCa and had digitized H&E slides, as well as whole-exome sequencing and bulk RNA-sequencing data.^30,31^ This cohort allowed for interrogation of molecular correlates of histologic features, but was not used for examination of clinical associations due to short median follow-up of 2.8 years and lack of information on DM. Additional information on cohort formation is listed in **Supp. Table 1**.

### Quantification of immune features from digitized H&E images

To understand the distribution of immune cells in our samples, we used CellViT,^32^ a deep learning model utilizing a pretrained vision transformer encoder, to identify immune cell nuclei^33^ for samples in the Discovery Cohort. Qualitatively, we observed that the spatial distribution of immune cells varied, from being scattered throughout tissue to forming dense clusters (**Supp. Fig. 1**). To quantify spatially dense immune clusters, we utilized density-based spatial clustering of applications with noise (DBSCAN),^34^ which is an unsupervised clustering algorithm that groups together points in dense regions (**Fig. 1a, Methods**). We then normalized the number of immune cells by the total number of cells (termed immune proportion) and the number of immune clusters by the amount of tissue. The median immune proportion was 4.3% (interquartile range [IQR] 3.0-5.9%), and the median immune cluster density was 2.4 per 25 μm^2^ of tissue (IQR 0-7.1). The density of immune clusters was moderately correlated with the immune proportion (R^2^=0.37). Notably, 30.9% (n=84) of samples did not have any immune clusters despite having immune proportions ranging from 1.1% to 8.5% (**Fig. 1b**). Similar results were observed for the Validation and TCGA Cohorts (**Supp. Fig. 2**). Representative whole slide images (WSI) of samples from the Discovery Cohort with variable values of immune proportion and immune cluster density are shown in **Fig. 1c**.

**Figure 1:**
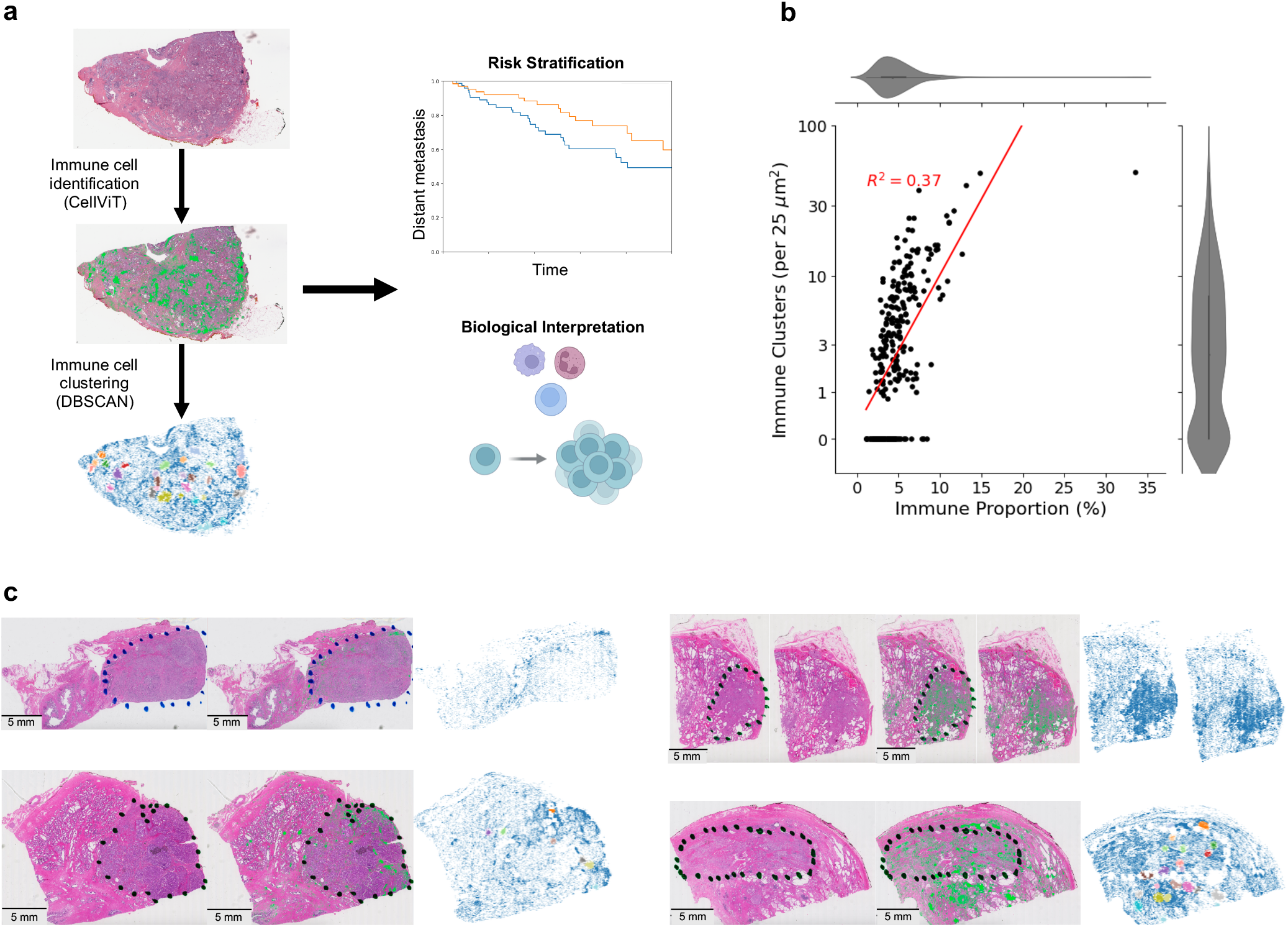
Framework for understanding the spatial architecture of immune cells. **a**, We applied a deep learning model (CellViT) to digitized H&E slides to identify immune cell locations, followed by spatial clustering using DBSCAN. This generated spatial immune representations that captured clinically and biologically meaningful features. **b**, Immune cluster density showed moderate correlation with immune proportion in the Discovery Cohort, but 30.8% of samples (n=84) lacked immune clusters despite variable amounts of immune cell presence. **c**, Representative examples from the Discovery Cohort illustrating low immune proportion without immune cluster (top left), high immune proportion without cluster (top right), low immune proportion with high cluster density (bottom left), and high immune proportion with high cluster density (bottom right).

### Association between histologic immune features and clinical outcomes

Given the ability to quantify interpretable features from the immune microenvironment of prostate tumors using WSIs, we investigated whether there was an association between immune histologic features and oncologic outcomes using Cox proportional hazards regression. We found that the immune proportion was not associated with DM, biochemical recurrence (BCR), or overall survival (OS) in the Discovery Cohort (P_int_≥0.20, **Supp. Table 2a**). Similar results were observed for the Validation Cohort (P_int_≥0.30, **Supp. Table 2b**).

Next, we investigated the association between immune cluster density and oncologic outcomes (**Fig. 2a, Table 2a**). In the Discovery Cohort, we observed that immune cluster density was associated with decreased risk of DM for Gleason 8-10 patients (adjusted hazard ratio [AHR] 0.42, 95% confidence interval [CI] 0.19-0.93) but not Gleason 6-7 patients (AHR 1.26, 95% CI 0.78-2.05), with a significant interaction (P_int_=0.019). We also observed a significant association between immune cluster density and OS for Gleason 8-10 patients (AHR 0.29, 95% CI 0.08-0.97) but not Gleason 6-7 patients (AHR 1.10, 95% CI 0.78-1.56; P_int_=0.038). Interestingly, no association between immune cluster density and BCR was observed for either Gleason 8-10 or 6-7 patients (P≥0.10).

**Figure 2:**
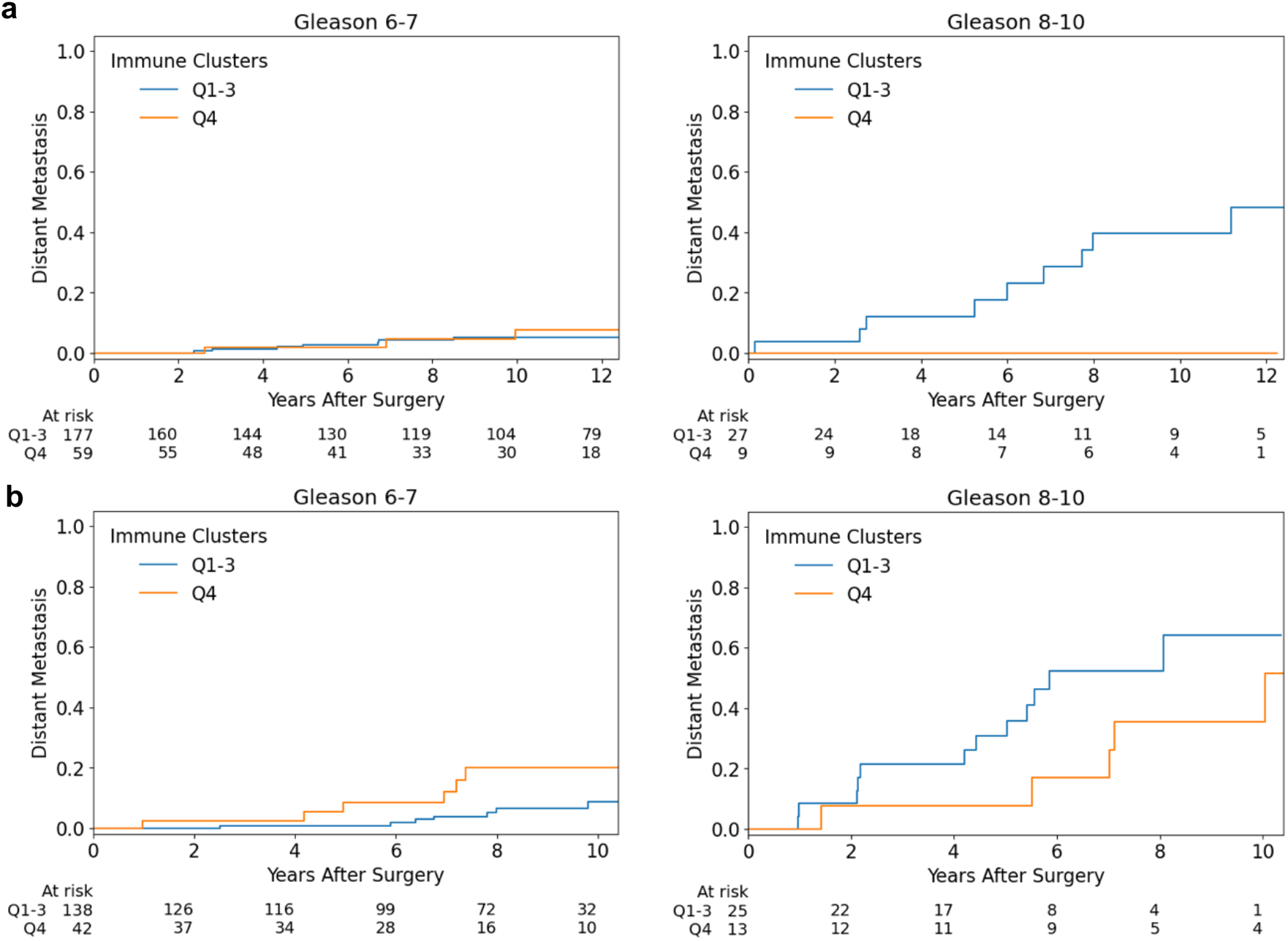
Cumulative incidence curves for distant metastasis. **a**,**b**, In both the Discovery (a) and Validation (b) Cohorts, patients were stratified by whether the immune cluster density fell within the top quartile (Q4 vs Q1-3). Immune cluster density did not distinguish outcomes in Gleason 6-7 disease but was prognostic in Gleason 8-10 disease.

**Table 1:** Baseline patient demographics. IQR: interquartile range, PSA: prostate-specific antigen, TCGA: The Cancer Genome Atlas.

|  | Discovery | Validation | TCGA |
| --- | --- | --- | --- |
| Cohort size | 272 | 218 | 326 |
| Age, median (IQR) | 63 (58-67) | 62 (57-65) | 61 (55-66) |
| Surgery year, median (IQR) | 1996 (1994-1997) | 2015 (2013-2017) | 2010 (2009-2011) |
| Gleason scores, n (%) | 6: 102 (38%)<br>7: 133 (49%)<br>8-10: 37 (14%) | 6: 23 (11%)<br>7: 157 (72%)<br>8-10: 38 (17%) | 6: 32 (10%)<br>7: 171 (52%)<br>8-10: 123 (38%) |
| pT stage, n (%) | T2: 209 (77%)<br>T3a: 45 (17%)<br>T3b/T4: 18 (7%) | T2: 128 (59%)<br>T3a: 59 (27%)<br>T3b/N1: 31 (14%) | T2: 122 (38%)<br>T3a: 108 (34%)<br>T3b/T4/N1: 92 (29%)<br>Missing: 4 (1%) |
| PSA, median (IQR) | 6.2 (4.5-9.2) | 5.9 (4.7-8.5) | Not available |
| Positive margins, n (%) | 94 (35%) | 57 (26%) | 91 (30%) |
| Median years of follow-up | 12.6 | 8.1 | 2.8 |

**Table 2:**
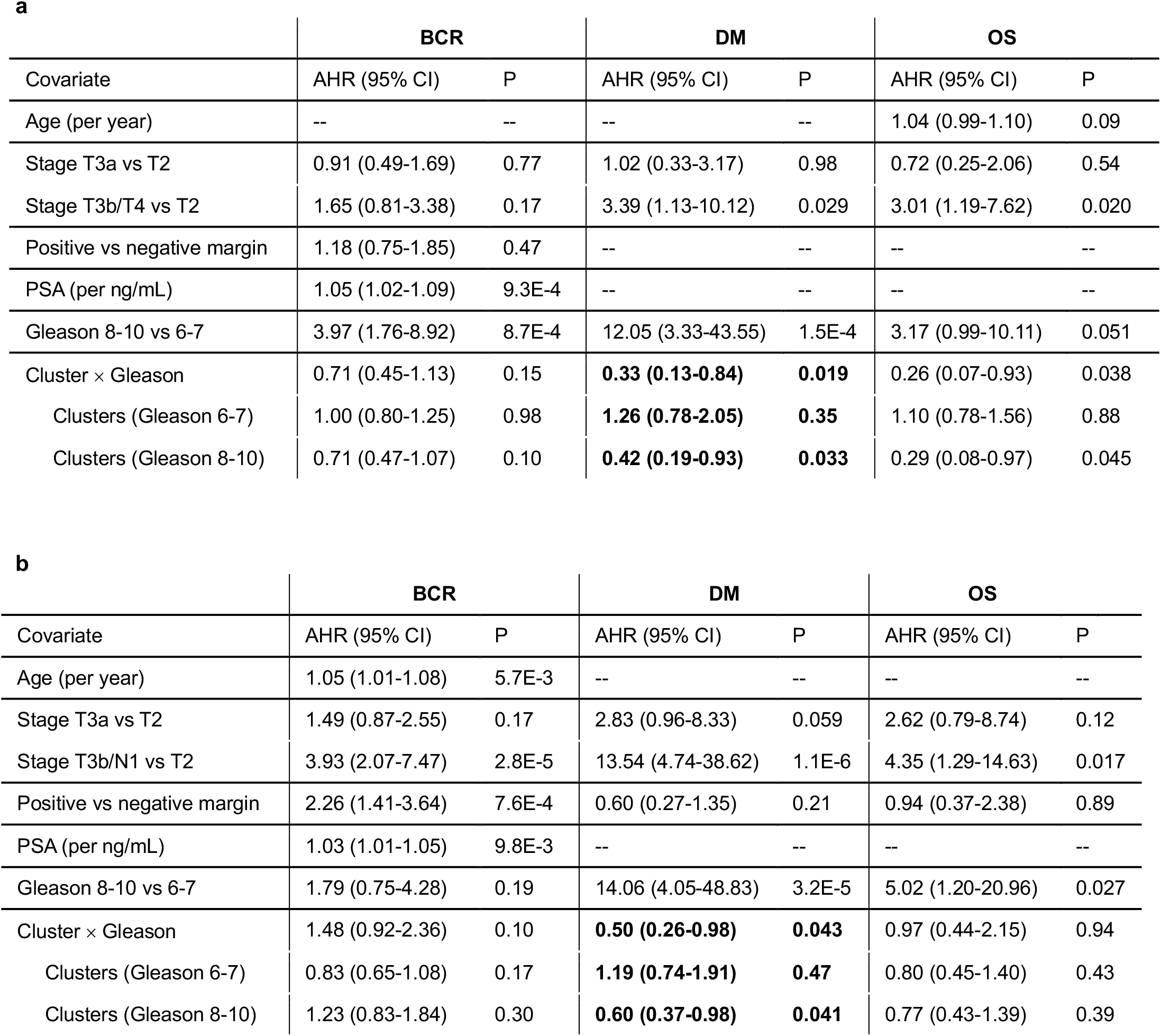
Cox proportional hazards regression models assessing effect of immune cluster density. **a**,**b**, In both the Discovery (a) and Validation (b) Cohorts, immune cluster density was associated with distant metastasis (DM) in Gleason 8-10 disease but not Gleason 6-7. In the Discovery Cohort (a), immune cluster density was associated with OS as well. However, no association with biochemical recurrence (BCR) was observed. AHR: adjusted hazard ratio, CI: confidence interval, PSA: prostate-specific survival.

In the Validation Cohort, we identified a similar association between immune cluster density and DM for Gleason 8-10 patients (AHR 0.60, 95% CI 0.37-0.98) but not Gleason 6-7 patients (AHR 1.19, 95% CI 0.74-1.91; P_int_=0.043; **Fig. 2b, Table 2b**). No significant association was observed between immune cluster density and either BCR (P≥0.10) or OS (P≥0.39).

Given the lack of an observed association between immune cluster density and BCR, despite an association with DM, we examined whether immune cluster density was associated with DM after development of BCR. We found that in the Discovery Cohort, immune cluster density was associated with different effects on DM after BCR (P_int_=0.036), with an AHR of 1.37 (95% CI 0.82-2.27) for Gleason 6-7 patients and 0.45 (95% CI 0.18-1.11) for Gleason 8-10 patients. Similar trends were observed in the Validation Cohort (**Supp. Table 3**).

### Genomic correlates of histologic immune cluster density

Having established a relationship between immune cell clusters in high-grade localized PCa and risk of DM, we then assessed whether specific genomic alterations were associated with immune cluster density using TCGA Cohort. Patients were stratified by Gleason score (6-7 vs 8-10), and within each group, we compared tumors in the top 10% of immune cluster density (high-cluster) to those in the bottom 90% (low-cluster). We did not detect differences in tumor mutational burden between low-cluster and high-cluster samples for either Gleason 6-7 (median 1.13 [IQR 0.90-1.53] vs 1.03 [IQR 1.00-1.30], P=0.85) or Gleason 8-10 (median 1.20 [IQR 0.97-1.47] vs 1.15 [IQR 0.99-1.37], P=0.59). We also did not detect differences in the fraction of genome altered^35^ for either Gleason 6-7 (median 4.2% [IQR 0.7-8.2%] vs 2.3% [0.2-6.3%], P=0.33) or Gleason 8-10 samples (median 12.1% [IQR 6.6-19.1%] vs 14.3% [2.8-27.3%], P=0.76; all by Wilcoxon rank-sum test). Additionally, there were no differences in the frequency of microsatellite instability, ETS fusions,^36^ or alterations in DNA damage repair genes,^37^ homologous recombination genes,^38^ PTEN,^39^ TP53,^40^ RB1, MYC, CHD1,^15^ or SPOP^41^ (all P≥0.12 by Fisher’s exact test), which have been previously linked to immune modulation (**Supp. Table 4**). Similar results were observed in the Validation Cohort where patients had data from panel sequencing, with the exception of PTEN in Gleason 6-7 disease (P=0.04), though this difference was not significant after multiple testing correction (**Supp. Table 5**), suggesting it is likely a spurious finding.

### Molecular immunologic correlates of histologic immune cluster density

Whereas we could not identify tumor-intrinsic genomic features linked to these machine learning-derived inferred microenvironmental patterns, we hypothesized that these patterns reflected underlying differences in immunologic states. To dissect immune cellular composition differences, we performed immune cell deconvolution using bulk RNA sequencing with CIBERSORTx,^42^ focusing on lymphoid, macrophage, and plasma cell populations.^15,19,43^ For Gleason 8-10 disease, high-cluster samples were proportionally enriched for CD8+ T cells (median 14.4% [IQR 8.9-19.2%] vs 9.9% [IQR 7.0-12.1%], P=0.023), activated memory CD4+ T cells (median 0% [IQR 0-3.5%] vs 0% [IQR 0-0%], P=0.014), and Tregs (median 2.3% [IQR 1.1-3.8%] vs 0.6% [IQR 0-1.9%], P=0.0036). For Gleason 6-7 samples, high-cluster samples were enriched for memory B cells (median 0% [IQR 0-1.8%] vs 0% [IQR 0-0%], P=0.0052) and γδ T cells (median 0% [IQR 0-0.2%] vs 0% [IQR 0-0%], P=0.038; all by Wilcoxon rank-sum test; **Supp. Fig. 3**).

Given the differential enrichment of B and T cell populations in high-cluster samples, we examined for evidence of clonal B and T cell expansions. We reconstructed BCR and TCR repertoires from bulk RNA sequencing using TRUST4 (**Methods**).^44^ A total of 209,792 BCR/TCR clonotypes were detected, but 32.7% (n=68,663) were singletons which may be nonfunctional and removed from downstream analysis (**Supp. Fig. 4** shows the clone count by receptor component including and excluding singletons). Clonality was then calculated, with values ranging from 0 (BCR/TCR repertoire characterized by even distribution of clonotypes) to 1 (BCR/TCR repertoire dominated by one or a few clonotypes).

When comparing low and high-cluster samples for either Gleason 6-7 or Gleason 8-10, differences IGH, IGK, or IGL clonality were not detected (P≥ 0.065). For TRB and TRA clonality, differences between low and high-cluster samples were not observed for Gleason 6-7 disease (P≥ 0.46) but were observed for Gleason 8-10 disease (P≤ 0.027; all by Wilcoxon rank-sum test). Overall, these results indicated increased clonal T cell populations, but not B cell populations, in Gleason 8-10 samples with high immune cluster density, and that this trend was not observed for Gleason 6-7 samples (**Fig. 3**). As microsatellite instability or high tumor mutational burden (defined as ≥10 nonsynonymous mutations/Mb) would be expected to be associated with clonal lymphocytic responses, these samples (n=3) were excluded. Similar results were also observed (**Supp. Fig. 5**), suggesting that the observed differences in T cell population clonality were not driven by hypermutated tumors.

**Figure 3:**
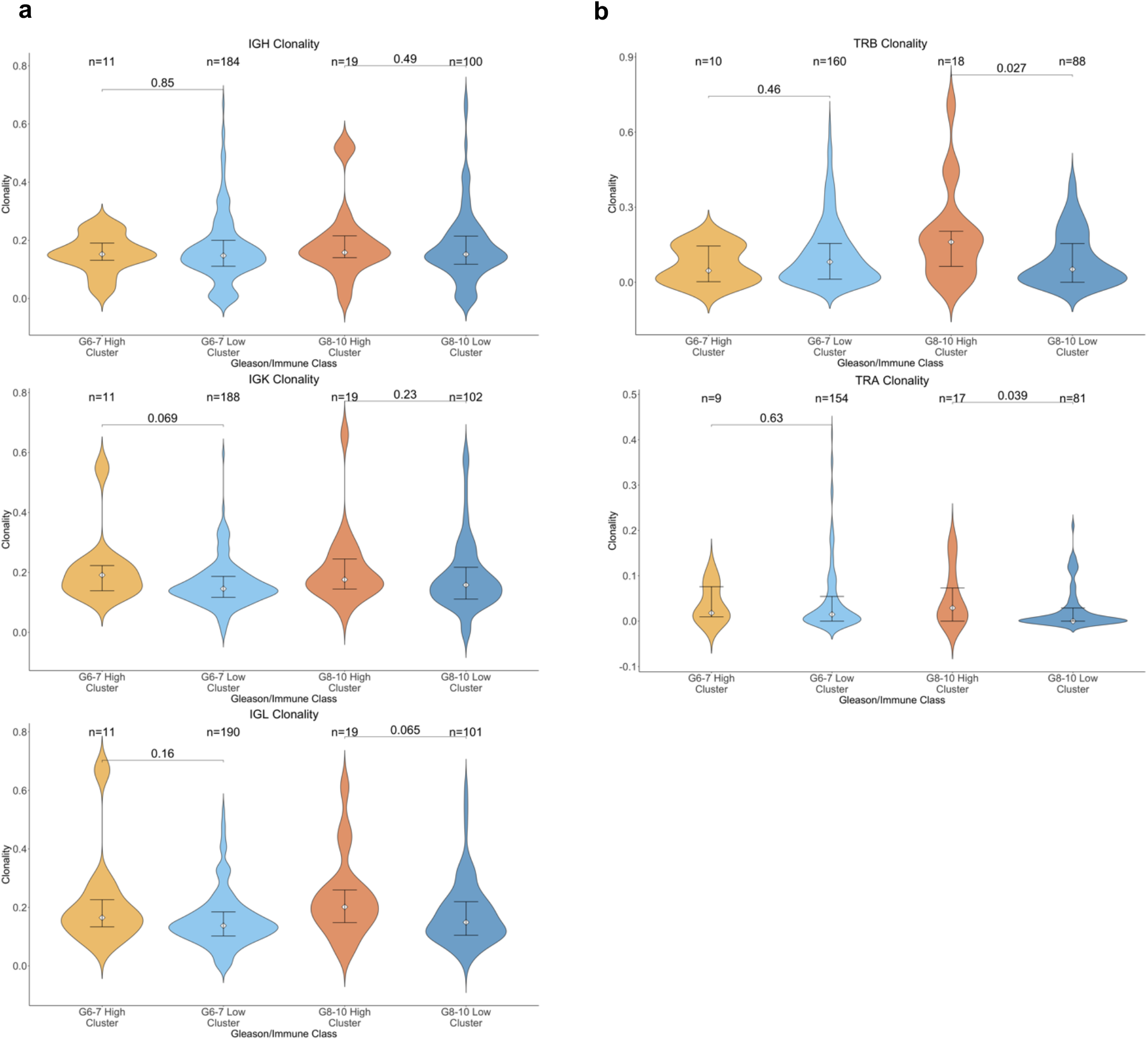
Clonality measurements from immune repertoire inference. **a**, BCR clonality were similar between high and low-cluster samples in both Gleason 6-7 and 8-10 disease. **b**, TCR clonality was higher in high-cluster samples compared to low-cluster samples in Gleason 8-10 disease, but not in Gleason 6-7 disease. Comparisons were made using Wilcoxon rank-sum test.

## DISCUSSION

The scarcity of robust biomarkers in localized PCa limits optimal risk stratification and personalized treatment recommendations. In this study, we identified and validated histologically defined spatial immune clustering as a novel computational pathology biomarker prognostic for distant metastasis in patients with high-grade PCa who undergo RP. Joint analysis of digitized H&E slides and sequencing data nominated T cell-mediated antitumor response as a potential mechanism underlying the observed clinical association.

This study has several potential implications. We demonstrate that immune spatial organization, rather than simple abundance, can offer independent prognostic value for DM in high-grade PCa. Notably, immune cluster density was not associated with BCR, which is of limited clinical utility.^45^ Rather, we showed that immune spatial organization can potentially discriminate high-risk BCR, where patients develop DM, from low-risk BCR. With established and emerging data supporting escalation of therapy for certain patients after RP (both in the adjuvant and salvage settings), including the addition of pelvic nodal radiation^46^ and increasing the duration^47^ and intensity^48,49^ of androgen receptor signaling blockade, immune cluster density may aid in personalizing treatment in this scenario.

Integrated analysis of histopathology and sequencing data implicated a T cell-mediated mechanism behind the observed protective effect against DM which is not associated with genomic alterations. High-cluster, high-grade samples were enriched for CD8+ T cells and activated memory CD4+ T cells and exhibited increased TCR clonality, consistent with a targeted antitumor immune response. The enrichment of Tregs may be indicative of local T cell exhaustion, but the geographic distribution of T cell states is not necessarily uniform;^50^ that is, local T cell exhaustion is not indicative of systemic T cell exhaustion. Future work will be needed to understand the drivers of this presumed antitumor immune response in select high-grade tumors and explore methods to modulate them.^51,52^

Several challenges and limitations of this study warrant consideration. First, the samples sizes of both the Discovery and Validation Cohorts are modest, though the prognostic value of immune cluster density was demonstrated in these two independent cohorts. Second, the shorter follow-up in the Validation Cohort, as well as advances in therapy leading to longer post-metastasis survival,^23^ may account for the absence of an association between immune cluster density and OS, in contrast to findings in the Discovery Cohort. Thus, further validation of our findings in additional, larger cohorts with robust follow-up periods is necessary prior to clinical adoption. Lastly, the immune cell identification model (CellViT) cannot differentiate between different immune cell subtypes (e.g., B vs T cells) or states (e.g., exhausted vs not) and may explain the lack of an observed association between immune proportion and clinical outcomes. As such, we focused on immune spatial architecture, but the cellular compositions of these clusters remain unknown though could be revealed through utilization of emerging spatial profiling techniques.^53^

Taken together, our findings highlight immune spatial architecture as a clinically relevant and biologically informed biomarker that may guide prognostication and future therapeutic strategies in high-grade PCa. The presence of clonal T cell populations in high-cluster, high-grade tumors warrants further investigation to uncover the immunologic dynamics of this state. Broadly, biologically informed computational pathology approaches for biomarker discovery warrant further consideration for use in PCa.

## METHODS

### Patient cohorts and data acquisition

The Discovery Cohort consisted of patients who underwent RP at Massachusetts General Hospital. Clinical data were gathered through November 30, 2010. Representative slides were prepared as part of a prior study in approximately 2010^29^ and digitized using a Hamamatsu C9600-12 NanoZoomer 2.0 HT at 40 × (0.23 μm/pixel) between August and November 2023. The Validation Cohort consisted of patients who were seen at Brigham and Women’s Hospital/Dana-Farber Cancer Institute for their PCa care and enrolled onto the PROFILE research study. Follow-up data were gathered through March 20, 2025. Representative slides were prepared as part of the diagnostic surgical pathology from the RP and digitized using an Aperio ScanScope v1 at 20 × (0.50 μm/pixel) between October 2013 and February 2018. Key details of TCGA Cohort have been previously described.^30^ Slides were digitized at 40 × (0.25 μm/pixel) using an Aperio ScanScope v1 between 2011 through 2014.

### Sequencing analysis

For TCGA Cohort, whole-exome sequencing was previously uniformly processed,^31^ and finalized calls for somatic single nucleotide variants, insertions/deletions, copy number alterations, and structural variants were used. For tumor suppressors, genomic alterations of interest included insertions/deletions, nonsense mutations, splice region alterations, translational start site alterations, deep deletions, and pathogenic missense mutations in COSMIC v98.^54^ For oncogenes, genomic alterations of interest included high amplifications and pathogenic missense mutations in COSMIC. Data on MSISensor score and fraction genome altered were obtained from cBioPortal on August 18, 2023,^55^ and microsatellite instability was determined based on MSISensor score >3.5.^56^ DNA damage repair genes consisted of BRCA1, BRCA2, ATM, BARD1, BRIP1, CDK12, CHEK1, CHEK2, FANCA, FANCL, NBN, PALB2, RAD51, RAD51B, RAD51C, RAD51D, and RAD54L.^57^ Homologous recombination genes consisted of BRCA1, BRCA2, and ATM.

For the bulk RNA sequencing, FASTQ files were used for TRUST4. Specifically, adapters (“AGATCGGAAGAGCACACGTCTGAACTCCAGTCA” and “AGATCGGAAGAGCGTCGTGTAGGGAAAGAGTGT”) were first trimmed using cutadapt v2.2.^58^ Next, TRUST4 v1.0.4^44^ was utilized, with the gene reference being the HG19 BCR/TCR reference and the gene annotation being IMGT, both included in the TRUST4 package. Clonality was then calculated as: 

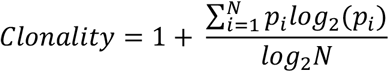

where *N* is the number of unique clonotypes and *p*_*i*_ is the frequency of clonotype *i*. Samples with 0-1 clonotypes cannot have clonality calculated and removed from comparisons.

Additionally, the FASTQ files without the adapters underwent two-pass STAR alignment.^59^ Samples with <48 base pair read lengths and <75% unique mapping reads were removed. Next, RNA expression was quantified using RSEM^60^ with a gencode30 gene annotation reference. Transcripts per million (TPM) measurements were calculated and used to impute cell fractions with the web-based implementation of CIBERSORTx^42^ using the LM22 gene signature matrix (performed on February 27, 2025).

For the Validation Cohort, patients underwent panel sequencing which provided data on single nucleotide variants, insertions/deletions, and copy number alterations.^61^ Only genes covered in all versions of the panel were analyzed; this excluded the analysis of CHD1, SPOP, ETS fusion, and certain genes in the DNA damage repair group (BARD1, CDK12, CHEK1, FANCL, RAD51, RAD51B, RAD51C, RAD51D, and RAD54L). Tumor mutational burden data was not available for the majority of the cohort, including all Gleason 8-10 patients, and hence not analyzed.

### Image analysis workflow

Images underwent quality control using GrandQC,^62^ a deep learning model which can segment and classify various artifacts, including air/edge, pen marking, out-of-focus, fold, and dark spot/foreign. We utilized the 10× model checkpoint for GrandQC v1, accessed on December 21, 2024. In reviewing GrandQC outputs, we noted that the model often labels pen marking as dark spot/foreign. As such, we considered artifact areas to be air/edge, out-of-focus, and fold, and we excluded samples with ≥ 5% artifact by area to ensure the analysis of high-quality slides. GrandQC also provided tissue segmentations, which were used to calculate the amount of tissue area. Next, we inferred immune cell nucleus segmentations using CellViT (accessed on June 13, 2024) with the recommended settings of 1024 × 1024-pixel patches and 64-pixel overlap on the highest magnification available for each slide (i.e., 40 × for Discovery and TCGA, 20 × for Validation). CellViT has demonstrated improved performance over prior models for cell nuclei instance segmentation and classification.^32^ CellViT was trained on nearly 200,000 nuclei of 5 classes (neoplastic, non-neoplastic epithelial, inflammatory/immune, connective, and dead) from 19 cancer types, and the immune class consists of lymphoid and macrophage cells.^33^ The encoder was a vision transformer (ViT-S) pretrained on 10,678 whole-slide H&E-stained images.^63^ The location of the immune cell was calculated based on the centroid of the nucleus segmentation. In TCGA where some patients had >1 slide, only the slide with the highest immune proportion was kept for downstream analysis. Approximately 120 representative WSIs with immune cell and immune cluster annotations, enriched for high-grade samples and spanning all three cohorts, were reviewed by an experienced pathologist (A.A.) to ensure that annotations were high-quality.

Next, we utilized DBSCAN,^34^ from the Python *sklearn* implementation, to identify immune spatial clusters. DBSCAN requires two parameters, *eps* (maximum neighborhood radius) and *min_samples* (the minimum number of nuclei to form a cluster), which were determined by applying the following approach to the Discovery Cohort. In Step 1, we evaluated 5 potential *eps* values (500, 750, 1000, 1250, 1500 pixels; 1 pixel=0.23 μm) and 4 *min_samples* values (250, 500, 750, 1000 nuclei). Biologically appropriate candidate *eps* values were chosen based on the approximate size of tertiary lymphoid structures^64^ while *min_samples* values were informed by visual inspection of representative slides. For each parameter combination, we calculated immune cluster density normalized to tissue area (**Supp. Fig. 6**). We then prioritized *eps* values of 1000 and 1250 and *min_samples* values of 500, 750, and 1000 for Step 2, as these produced density distributions with low medians and pronounced right skew, consistent with prior findings that most localized prostate cancers are immune-depleted, with a subset showing immune enrichment.^17,18^ In Step 3, we performed Cox proportional hazards regressions between immune cluster density and DM (per below) and selected the parameter combination (*eps*=1250, *min_samples*=500) that yielded the lowest P-value for the interaction term in the model (**Supp. Fig. 7**).

### Survival analysis

We utilized Cox proportional hazards regression to examine the association between either immune proportion or immune cluster density (transformed as ln[density+1] due to data skewedness) and the primary outcome of DM and secondary outcome of BCR and OS, all calculated as time since RP. As additional exploratory analysis, we examined the association between immune cluster density and DM for patients who experienced a BCR, measured from time of BCR. For BCR and DM, death was treated as a censored event. Other potential covariates included age (years), PSA level at time of diagnosis, pathologic stage (T2, T3a, and T3b/T4 [Discovery] or T3b/N1 [Validation]), surgical margin status (positive and negative) and Gleason score from the RP specimen (Gleason 6-7 and 8-10). For the primary and secondary endpoints, we first performed univariable regression analysis. The multivariable model included immune proportion/immune cluster density; Gleason score; the interaction term between immune proportion/immune cluster density and Gleason score; as well as other covariates with P<0.10 on univariable regression. In the model exploring time between BCR and DM, we utilized the same covariates as the model exploring time from RP to DM. A small L2 regularization term (2.5E-3) was used in all multivariable models to reduce overfitting and enhance model generalizability.

For comparisons of sequencing analysis results, samples were divided into Gleason 6-7 vs Gleason 8-10. Within each Gleason group, we compared samples in the top 10% of immune cluster density (high-cluster) to those in the bottom 90% (low-cluster). The 10th percentile threshold was selected to enhance the detection of differences.

## Statistical analysis

All statistical analysis was performed using Python 3 (for image-related analysis) and R 4.4.1 (for genomics-related analysis). We compared categorical variables using two-sided Fisher’s exact test and continuous variables using two-sided Wilcoxon rank-sum test, utilizing the base *fisher*.*exact* and *wilcox*.*test* functions. Linear regression was performed using *scipy* in Python. Survival analysis including Cox proportional hazards regression and the plotting of Kaplan-Meier cumulative incidence curves was performed using *lifelines* in Python. Violin plots contained associated bar plots for 25^th^, 50^th^ (median), and 75^th^ quartiles. P<0.05 was considered statistically significant. This study complied with REMARK reporting guidelines for prognostic tumor biomarkers.^65^

## Acknowledgments

Our results are in part based upon data generated by the TCGA Research Network (https://www.cancer.gov/tcga). This work is supported by a Prostate Cancer Foundation Young Investigator Award (D.D.Y, K.S.), Prostate Cancer Foundation Challenge Award (E.M.V.A., M.E.T), SPORE P50CA272390 (E.M.V.A., C.L.W., D.D.Y., K.S.), P01CA228696 (E.M.V.A.), DOD W81XWH-21-PCRP-DSA (E.M.V.A.), and HT94252410415 (E.M.V.A.).

## Author contributions

D.D.Y. designed the experiments, performed data generation, analyzed and interpreted results, and drafted the original version of the manuscript. A.A. analyzed and interpreted results. J.W. interpreted results. A.J.H. interpreted results. A.A.B. performed data generation. E.S., J.P.C., and C.A.R. designed aspects of the bioinformatics analysis pipeline. J.P. interpreted results.

M.T.K. performed data generation. P.L.N. provided supervision and supported funding acquisition. T.K.C. interpreted results and supported funding acquisition. A.K.T. interpreted results. K.S. interpreted results and supported funding acquisition. M.E.T interpreted results and supported funding acquisition. C-L.W. performed data generation, interpreted results, provided supervision, and supported funding acquisition. E.M.V.A designed the experiments, interpreted results, provided supervision, and supported funding acquisition. All authors reviewed and edited versions of the manuscript.

## Data availability

TCGA digitized slides and bulk RNA sequencing data were downloaded from Genomic Data Commons. Processed whole exome sequencing data were obtained from a prior publication which uniformly processed the raw sequencing data.^31^ Clinical data for the Discovery Cohort are available from the prior publication.^29^ Digitized H&E images, sequencing data from the Validation Cohort, and clinical data from the Validation Cohort are available upon request, contingent on IRB approval and compliance with institutional policies regarding data security and use. Requests for academic, non-commercial use should be directed to the lead author via email. All requests will be reviewed in accordance with institutional and departmental guidelines to assess potential intellectual property or patient privacy constraints. Data sharing will require a formal Data Use Agreement. Additional information necessary to reproduce the analyses in this study is also available upon request from the lead contact.

## Code availability

A GitHub repository with code to reproduce the findings of this study will be posted once the manuscript has been published in a peer-reviewed journal.

