## Supplemental Tables and Figures for "Immune Spatial Organization Predicts Metastasis Risk in Aggressive Localized Prostate Cancer"

a

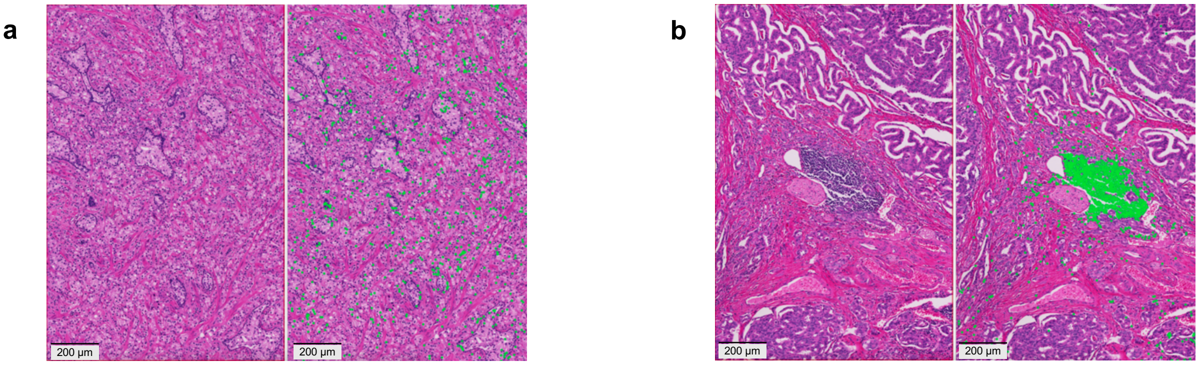


**Supplemental Figure 1: Representative examples of immune cell identification performed by CellViT.**
**a,b,** Sometimes immune cells are scattered throughout tissue (a; left is the H&E, and right is the H&E overlaid with green points for immune cell nuclei), while other times, they form dense clusters (b).

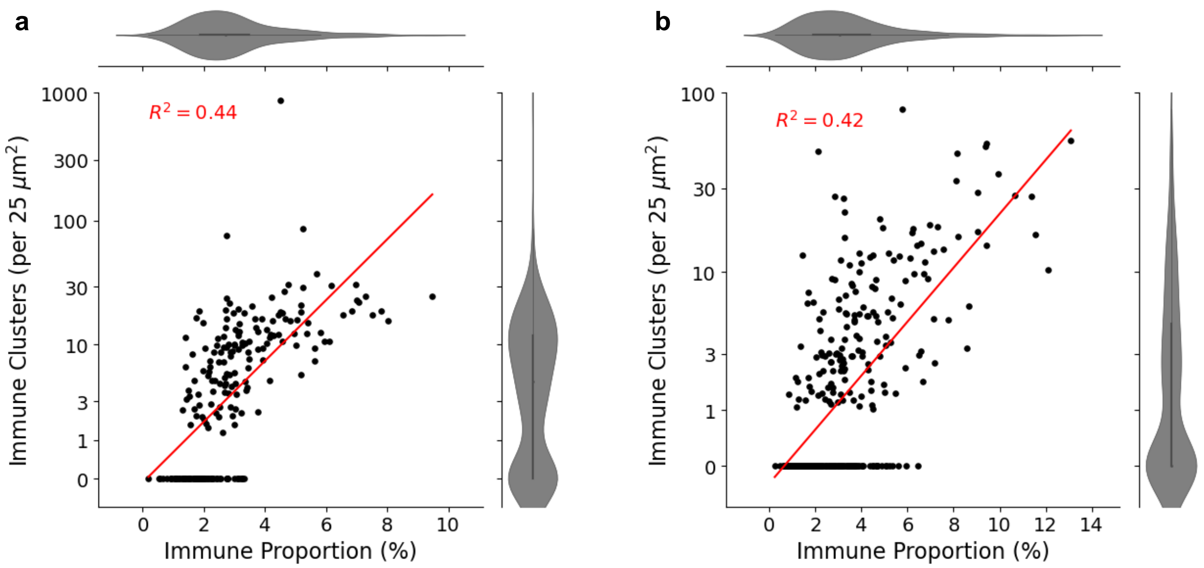


**Supplemental Figure 2: Scatter plots of immune cluster density vs immune proportion.**
**a,b,** Immune cluster density showed moderate correlation with immune proportion in the Validation (a) and TCGA (b) Cohorts, but 33.5% (n=73) and 50.6% (n=165), respectively, lacked immune clusters despite variable amounts of immune cell presence.

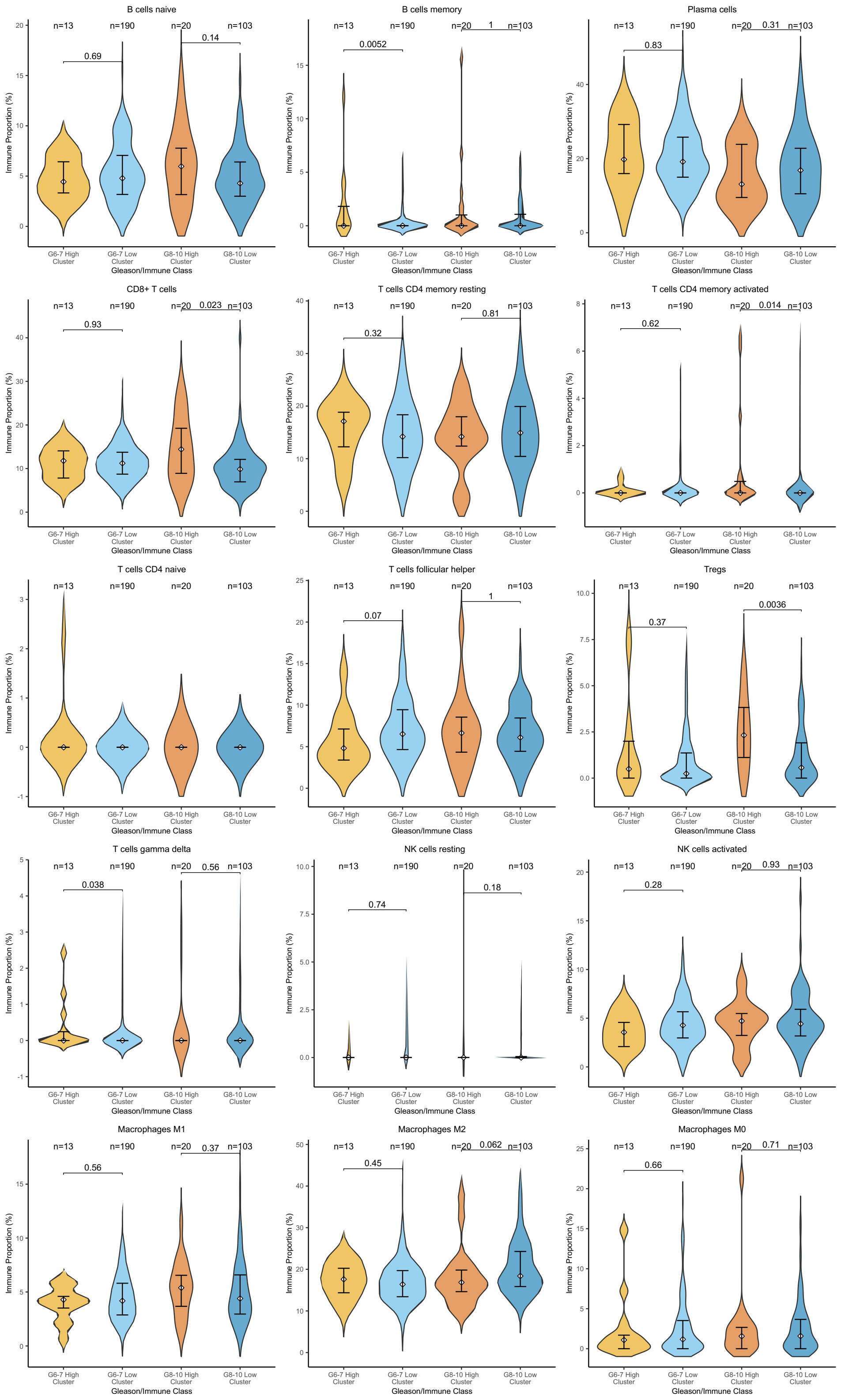

**Supplemental Figure 3: Immune cell type deconvolution using CIBERSORTx for lymphoid, macrophage, and plasma cell populations.**
Comparisons were made using Wilcoxon rank-sum test.

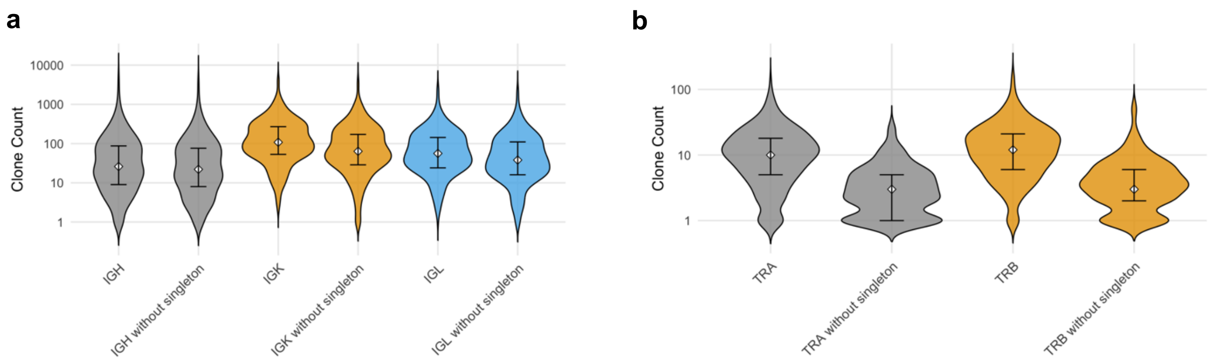


**Supplemental Figure 4: Clonotype count with and without singletons.**
**a,b,** The removal of singletons did not clearly impact the distribution of clonotype counts for BCR components (a) but did for TCR components (b), suggesting that TCR singletons are unlikely to be biologically pertinent. As such, they were removed from clonality measurements.

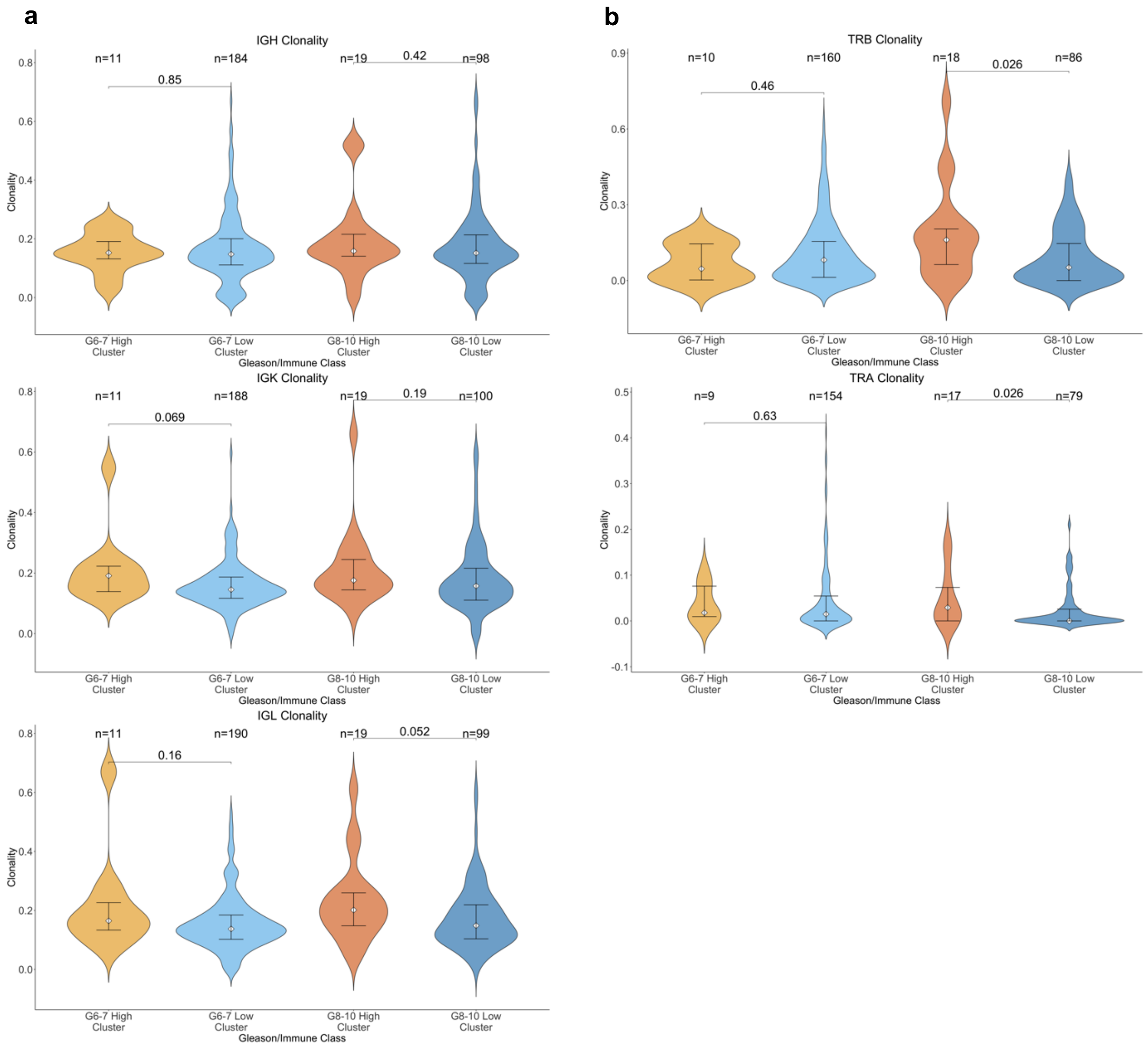


**Supplemental Figure 5: Clonality measurements from immune repertoire inference after removing the three samples with microsatellite instability.**
**a,** BCR components were similar between high and low-cluster samples in both Gleason 6-7 and 8-10 disease. **b,** TCR clonality was higher in high-cluster samples compared to low-cluster samples in Gleason 8-10 disease, but not in Gleason 6-7. Patients with 0-1 clonotypes were excluded from clonality analysis. Comparisons were made using Wilcoxon rank-sum test.

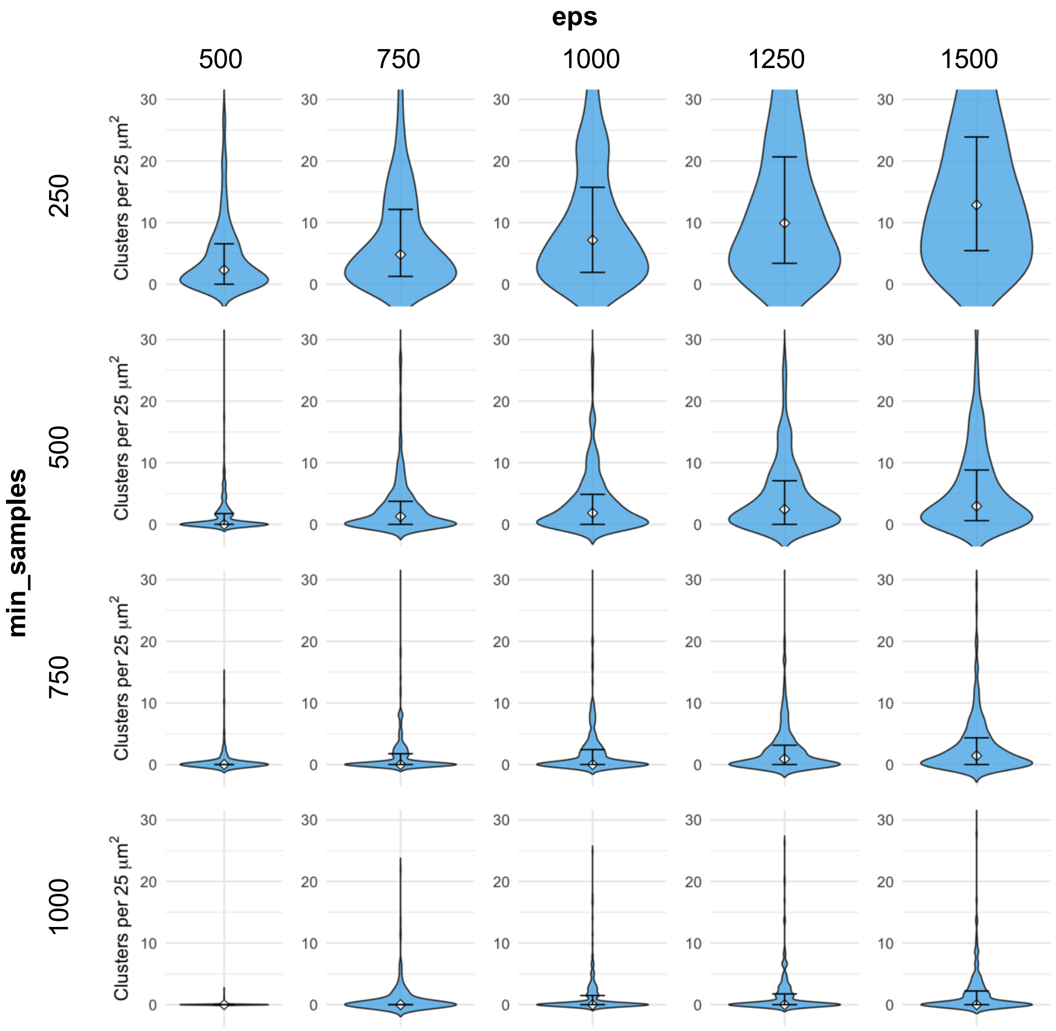


**Supplemental Figure 6: Immune cluster density using different parameter combinations for DBSCAN.**

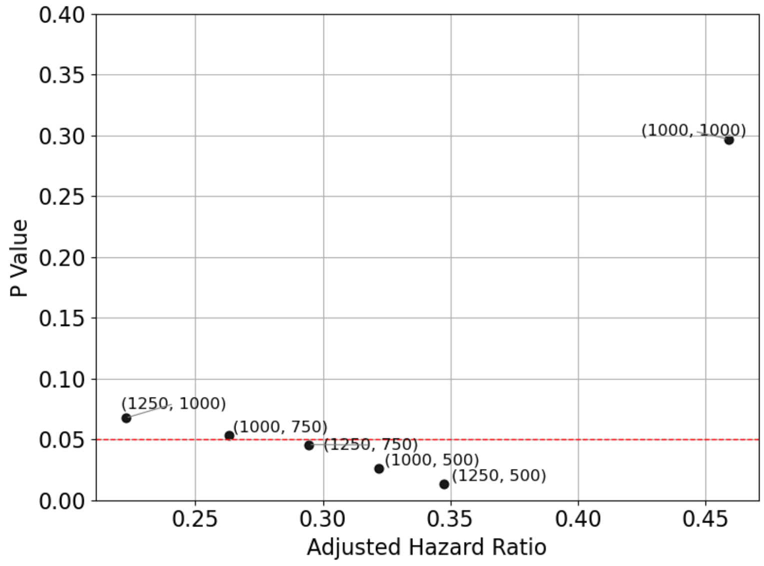


**Supplemental Figure 7: Scatter plot of P-values and adjusted hazard ratios for the Cox regression model for time to distant metastasis, using different combinations of DBSCAN parameters for determining immune cluster density.**
Each label consists of (*eps*, *min_samples*). The combination of *eps* = 1250 and *min_samples* = 500 produced the lowest P-value. Red line: P=0.05.

| Exclusion reason | Discovery (n=450) | Validation (n=258) | TCGA (n=394) |
|---|---|---|---|
| Missing baseline characteristic data (PSA, pT stage, Gleason score, or margin status) | 60 | 16 | N/A |
| Did not pass slide quality control | 118 | 24 | 34 |
| RNA sequencing did not pass QC or not accessible | N/A | N/A | 34 |
| Final cohort | 272 | 218 | 326 |

**Supplemental Table 1: Patient cohort formation.**
PSA: prostate-specific antigen, QC: quality control.

**a**

| | BCR | | DM | | OS | |
|---|---|---|---|---|---|---|
| Covariate | AHR (95% CI) | P | AHR (95% CI) | P | AHR (95% CI) | P |
| Age (per year) | -- | -- | -- | -- | 1.05 (1.00-1.10) | 0.06 |
| Stage T3a vs T2 | 0.88 (0.48-1.62) | 0.69 | 1.28 (0.42-3.90) | 0.67 | 0.77 (0.27-2.19) | 0.63 |
| Stage T3b/T4 vs T2 | 1.87 (0.95-3.70) | 0.07 | 3.41 (1.21-9.62) | 0.021 | 2.68 (1.07-6.74) | 0.036 |
| Positive vs negative margin | 1.27 (0.82-1.98) | 0.29 | -- | -- | -- | -- |
| PSA (per ng/mL) | 1.05 (1.02-1.09) | 1.8E-3 | -- | -- | -- | -- |
| Gleason 8-10 vs 6-7 | 1.60 (0.70-3.65) | 0.27 | 6.90 (1.26-37.76) | 0.026 | 1.44 (0.24-8.62) | 0.69 |
| **Immune proportion × Gleason** | **1.08 (0.96-1.23)** | **0.20** | **0.87 (0.61-1.24)** | **0.45** | **0.94 (0.64-1.37)** | **0.73** |
| **Immune (Gleason 6-7)** | **0.98 (0.89-1.09)** | **0.74** | **0.94 (0.75-1.19)** | **0.62** | **0.93 (0.78-1.11)** | **0.41** |
| **Immune (Gleason 8-10)** | **1.07 (0.99-1.15)** | **0.09** | **0.82 (0.61-1.11)** | **0.20** | **0.87 (0.61-1.24)** | **0.44** |

**b**

| | BCR | | DM | | OS | |
|---|---|---|---|---|---|---|
| Covariate | AHR (95% CI) | P | AHR (95% CI) | P | AHR (95% CI) | P |
| Age (per year) | 1.04 (1.01-1.08) | 9.9E-3 | -- | -- | -- | -- |
| Stage T3a vs T2 | 1.45 (0.84-2.49) | 0.18 | 2.91 (0.99-8.56) | 0.054 | 2.81 (0.85-9.33) | 0.09 |
| Stage T3b/N1 vs T2 | 3.93 (2.10-7.36) | 1.9E-5 | 13.38 (4.66-38.39) | 1.4E-6 | 4.35 (1.31-14.52) | 0.017 |
| Positive vs negative margin | 2.32 (1.45-3.71) | 4.5E-4 | 0.56 (0.25-1.28) | 0.17 | 0.99 (0.39-2.50) | 0.98 |
| PSA (per ng/mL) | 1.04 (1.01-1.06) | 6.4E-4 | -- | -- | -- | -- |
| Gleason 8-10 vs 6-7 | 2.99 (1.19-7.47) | 0.019 | 9.89 (2.26-43.21) | 2.3E-3 | 2.44 (0.45-13.39) | 0.30 |
| **Immune proportion × Gleason** | **1.02 (0.80-1.30)** | **0.87** | **0.81 (0.55-1.20)** | **0.30** | **1.27 (0.75-2.16)** | **0.37** |
| **Immune (Gleason 6-7)** | **0.93 (0.77-1.11)** | **0.41** | **1.02 (0.76-1.38)** | **0.89** | **0.70 (0.44-1.11)** | **0.13** |
| **Immune (Gleason 8-10)** | **0.95 (0.80-1.12)** | **0.51** | **0.83 (0.64-1.08)** | **0.16** | **0.89 (0.67-1.19)** | **0.44** |

**Supplemental Table 2: Cox proportional hazards regression models assessing effect of immune proportion.**
**a,b,** Immune proportion was not associated with biochemical recurrence (BCR), distant metastasis (DM), overall survival (OS) in either the Discovery (a) or Validation Cohorts (b). AHR: adjusted hazard ratio, CI: confidence interval, PSA: prostate-specific survival.

| | Discovery | | Validation | |
|---|---|---|---|---|
| Covariate | AHR (95% CI) | P | AHR (95% CI) | P |
| Age (per year) | -- | -- | -- | -- |
| Stage T3a vs T2 | 0.63 (0.19-2.07) | 0.45 | 1.54 (0.51-4.61) | 0.44 |
| Stage T3b/T4 vs T2 | 1.50 (0.53-4.25) | 0.44 | 3.87 (1.37-10.91) | 0.011 |
| Positive vs negative margin | -- | -- | 0.56 (0.26-1.21) | 0.14 |
| PSA (per ng/mL) | -- | -- | -- | -- |
| Gleason 8-10 vs 6-7 | 4.68 (1.32-16.59) | 0.017 | 6.09 (1.73-21.46) | 4.9E-3 |
| **Cluster × Gleason** | **0.33 (0.12-0.93)** | **0.036** | **0.56 (0.29-1.09)** | **0.089** |
| **Clusters (Gleason 6-7)** | **1.37 (0.82-2.27)** | **0.23** | **1.11 (0.71-1.75)** | **0.64** |
| **Clusters (Gleason 8-10)** | **0.45 (0.18-1.11)** | **0.084** | **0.62 (0.37-1.05)** | **0.077** |

**Supplemental Table 3: Cox proportional hazards regression models assessing time from biochemical recurrence (BCR) to distant metastasis (DM).**
Immune cluster density was associated with different effects on DM after BCR by Gleason score in the Discovery Cohort. A similar trend was observed in the Validation Cohort. AHR: adjusted hazard ratio, CI: confidence interval, PSA: prostate-specific survival.

| Gleason 8-10 (n=123) | Cluster density | Not altered | Altered | P |
|---|---|---|---|---|
| MSI | Low | 101 (98.1%) | 2 (1.9%) | 0.42 |
| | High | 19 (95.0%) | 1 (5.0%) | |
| DDR | Low | 89 (86.4) | 14 (13.6%) | 0.12 |
| | High | 20 (100%) | 0 (0%) | |
| HRD | Low | 98 (95.1%) | 5 (4.9%) | 0.59 |
| | High | 20 (100%) | 0 (0%) | |
| PTEN | Low | 89 (86.4%) | 14 (13.6%) | 1 |
| | High | 17 (85.0%) | 3 (15.0%) | |
| TP53 | Low | 80 (77.7%) | 23 (22.3%) | 0.56 |
| | High | 14 (70.0%) | 6 (30.0%) | |
| RB1 | Low | 97 (94.2%) | 6 (5.8) | 1 |
| | High | 19 (95.0%) | 1 (5.0) | |
| MYC | Low | 103 (94.2%) | 6 (5.8%) | 1 |
| | High | 19 (95.0%) | 1 (5.0%) | |
| CHD1 | Low | 94 (91.3%) | 9 (8.7%) | 1 |
| | High | 18 (90.0%) | 2 (10.0%) | |
| SPOP | Low | 95 (92.2%) | 8 (7.8%) | 0.67 |
| | High | 18 (90.0%) | 2 (10.0%) | |
| ETS fusion | Low | 46 (44.7%) | 57 (55.3%) | 0.23 |
| | High | 12 (60.0%) | 8 (40.0%) | |

| Gleason 6-7 (n=203) | Cluster density | Not altered | Altered | P |
|---|---|---|---|---|
| MSI | Low | 190 (100%) | 0 (0%) | 1 |
| | High | 13 (100%) | 0 (0%) | |
| DDR | Low | 163 (85.8%) | 27 (14.2%) | 1 |
| | High | 12 (92.3%) | 1 (7.7%) | |
| HRD | Low | 181 (95.3%) | 9 (4.7%) | 1 |
| | High | 13 (100%) | 0 (0%) | |
| PTEN | Low | 178 (93.7%) | 12 (6.3%) | 1 |
| | High | 13 (100%) | 0 (0%) | |
| TP53 | Low | 179 (94.2%) | 11 (5.8%) | 0.56 |
| | High | 12 (92.3%) | 1 (7.7%) | |
| RB1 | Low | 185 (97.4%) | 5 (2.6%) | 1 |
| | High | 13 (100%) | 0 (0%) | |
| MYC | Low | 185 (97.4%) | 5 (2.6%) | 1 |
| | High | 13 (100%) | 0 (0%) | |
| CHD1 | Low | 172 (90.5%) | 18 (9.5%) | 0.61 |
| | High | 13 (100%) | 0 (0%) | |
| SPOP | Low | 174 (91.6%) | 16 (8.4%) | 0.60 |
| | High | 13 (100%) | 0 (0%) | |
| ETS fusion | Low | 72 (37.9%) | 118 (62.1%) | 1 |
| | High | 5 (38.5%) | 8 (61.5%) | |

**Supplemental Table 4: Prevalence of genomic alterations between low and high-cluster samples for both Gleason 6-7 and 8-10 disease in TCGA Cohort.**

Comparisons were made using Fisher's exact test. DDR: DNA damage repair, HRD: homologous recombination, MSI: microsatellite instability.

| Gleason 8-10 (n=37) | Cluster density | Not altered | Altered | P |
|---|---|---|---|---|
| MSI | Low | 7 (100%) | 0 (0%) | N/A |
| | High | 30 (100%) | 0 (0%) | |
| DDR | Low | 28 (93.3%) | 2 (6.7%) | 1 |
| | High | 7 (100%) | 0 (0%) | |
| HRD | Low | 29 (96.7%) | 1 (3.3%) | 1 |
| | High | 7 (100%) | 0 (0%) | |
| PTEN | Low | 27 (90.0%) | 3 (10.0%) | 1 |
| | High | 7 (100%) | 0 (0%) | |
| TP53 | Low | 24 (80.0%) | 6 (20.0%) | 1 |
| | High | 6 (85.7%) | 1 (14.3%) | |
| RB1 | Low | 30 (100%) | 0 (0%) | N/A |
| | High | 7 (100%) | 0 (0%) | |
| MYC | Low | 30 (100%) | 0 (0%) | N/A |
| | High | 7 (100%) | 0 (0%) | |

| Gleason 6-7 (n=177) | Cluster density | Not altered | Altered | P |
|---|---|---|---|---|
| MSI | Low | 162 (100%) | 0 (0%) | N/A |
| | High | 15 (100%) | 0 (0%) | |
| DDR | Low | 158 (97.5%) | 4 (2.5%) | 0.36 |
| | High | 14 (93.3%) | 1 (6.7%) | |
| HRD | Low | 159 (98.1%) | 3 (1.9%) | 0.30 |
| | High | 14 (93.3%) | 1 (6.7%) | |
| PTEN | Low | 155 (95.7%) | 7 (4.3%) | 0.04 |
| | High | 12 (80.0%) | 3 (20.0%) | |
| TP53 | Low | 155 (95.7%) | 7 (4.3%) | 1 |
| | High | 15 (100%) | 0 (0%) | |
| RB1 | Low | 162 (100%) | 0 (0%) | N/A |
| | High | 15 (100%) | 0 (0%) | |
| MYC | Low | 162 (100%) | 0 (0%) | N/A |
| | High | 15 (100%) | 0 (0%) | |

**Supplemental Table 5: Prevalence of genomic alterations between low and high-cluster samples for both Gleason 6-7 and 8-10 disease in the Validation Cohort.**
Comparisons were made using Fisher's exact test. DDR: DNA damage repair, HRD: homologous recombination, MSI: microsatellite instability.
